# The vP300 Framework: Trial-to-Trial Variability in the P300 Event-Related Potential as a Signal of Locus Coeruleus-Norepinephrine Dynamics

**DOI:** 10.64898/2026.07.30.741927

**Authors:** Jianhua Su, Huilin Hu, Xin Su, Yannan Huang

**Affiliations:** School of Foreign Languages, Yang-En University, Quanzhou, China; College of Foreign Languages, Fujian Normal University, Fuzhou, China; La Trobe Business School, La Trobe University, Melbourne, Victoria, Australia; Department of Foreign Languages, Jimei University Chengyi College, Xiamen, China

**Keywords:** P300, event-related potential, trial-to-trial variability, coefficient of variation, locus coeruleus-norepinephrine, individual differences, ERP CORE, single-trial analysis, simulation methods, open science, adaptive gain, cognitive flexibility

## Abstract

The P300 event-related potential is one of the most studied electrophysiological markers of cognition, yet trial-to-trial variability within conditions has traditionally been treated as measurement error.The vP300 framework proposes that P300 variability, quantified by the coefficient of variation (CV), may reflect locus coeruleus-norepinephrine (LC-NE) mode dynamics. A computational simulation (N = 80) tested three hypotheses. Empirical P3 oddball data from the ERP CORE dataset (N = 39) provided partial validation. Mean amplitude and CV were orthogonal in simulation (r = −0.14, n.s.) and moderately correlated in empirical data (r = −0.418, p = .008), with 82.5% of CV variance independent of mean amplitude. CV significantly predicted cognitive flexibility (r = −0.319, p = .004) while mean amplitude did not (r = .010, p = .928). Trial-to-trial P300 variability carries information orthogonal to traditional mean amplitude, supporting a potential revision of how variability is treated in ERP research.

## Introduction

The P300 event-related potential (ERP) is one of the most studied electrophysiological markers of human cognition. Discovered by Sutton et al. [1] and linked to attentional allocation and context updating [2,3], the P300 has been a cornerstone of cognitive neuroscience for five decades. However, a fundamental methodological assumption has constrained progress: trial-to-trial variability within a condition is treated as random measurement error [4]. The consequence is that ERP researchers systematically discard the very information that may differentiate individuals with similar mean amplitudes but fundamentally different cognitive profiles. This assumption not only limits the explanatory power of ERP measures but may explain why P300 amplitude correlates inconsistently with behavior across studies despite robust within-study effects.

Several lines of evidence demonstrate that trial-to-trial variability is structured rather than random. Arieli et al. [7] showed that ongoing neural activity accounts for substantial trial-to-trial variance in evoked cortical responses. Iemi et al. [6] demonstrated that pre-stimulus alpha power systematically predicts single-trial ERP amplitude, and sequential effects documented over four decades ago [8,16] show that P300 amplitude depends on preceding trial history. However, these findings have not translated into a practical analytical framework because: (1) averaging continues to be the default analytical approach in ERP research, masking trial-to-trial dynamics; (2) no established metric exists for quantifying individual differences in trial-to-trial variability; and (3) the existing evidence, while suggestive of structured variability, lacks a unified theoretical framework linking variability to a specific neural system with testable predictions.

The locus coeruleus-norepinephrine (LC-NE) system provides a plausible neural basis for structured trial-to-trial P300 variability. The adaptive gain theory [11] distinguishes between phasic LC-NE mode (focused attention, consistent responses, low variability) and tonic mode (exploratory behavior, flexible responses, high variability). The P300 depends critically on LC-NE integrity: LC lesions in animals abolish the P300, and noradrenergic drugs systematically modulate P300 amplitude [12,13]. If LC-NE mode fluctuates from trial to trial, P300 amplitude variability should directly index the range and flexibility of LC-NE dynamics.

The vP300 framework directly addresses these gaps by proposing that the coefficient of variation (CV = SD / mean x 100) of P300 amplitude across trials operationalizes LC-NE mode dynamics. Unlike prior approaches, CV is: (1) computable from any existing averaged P300 dataset without requiring new data collection, (2) quantifiable per participant, enabling individual differences research that mean amplitude alone cannot capture, and (3) grounded in adaptive gain theory, generating specific, testable predictions about the relationship between P300 variability and cognitive function. We test three hypotheses:

H1: P300 mean amplitude and P300 CV are orthogonal dimensions of individual differences;

H2: P300 CV predicts cognitive flexibility while mean amplitude does not;

H3: trial-by-trial P300 amplitude sequences exhibit structured (1/f-like) spectral characteristics consistent with temporally correlated LC-NE dynamics rather than white noise.

The vP300 framework has the potential to unlock a new dimension of analysis for thousands of existing P300 datasets collected over five decades, potentially revising both the methodological approach and theoretical interpretation of one of the most studied ERP components in cognitive neuroscience. To test these hypotheses under conditions with known ground truth parameters - a necessary step for validating a new analytical framework - we employed a computational simulation approach. As noted by Cohen [5], simulation-based validation is essential for establishing the internal validity of novel analytical frameworks prior to their application to empirical data.

### The P300 Component and the Oddball Paradigm

The P300 was first described by Sutton et al. [1], who observed that unpredictable stimuli elicited substantially larger late-positive potentials than predictable stimuli. This seminal discovery established the oddball paradigm as the primary experimental tool for studying the P300 component and opened a new field of cognitive electrophysiology. Donchin [2] formalized the context-updating theory, which proposes that the P300 reflects the active updating of working memory representations when the brain detects a mismatch between expected and actual sensory input. This theory remains the dominant theoretical framework for understanding the P300 after more than four decades, supported by extensive evidence demonstrating that P300 amplitude is inversely related to stimulus probability and directly related to task relevance. The theory has been refined and elaborated by Polich [3], who demonstrated that the P300 can be fractionated into the P3a (frontocentral, novelty-driven, reflecting involuntary attentional orienting) and the P3b (centroparietal, task-relevant, reflecting active context updating).

Neural generators of the P300 have been identified through converging evidence from source localization, intracranial recording, and neuroimaging studies. The primary sources include the temporoparietal junction, the medial temporal lobe including the hippocampus and parahippocampal regions, the prefrontal cortex, and critically, the locus coeruleus [12]. This distributed network reflects the involvement of multiple cognitive processes, including attentional orienting, memory encoding and retrieval, and noradrenergic modulation of cortical processing. The involvement of the LC is particularly important for the present investigation, as it provides the anatomical and physiological basis for linking P300 variability to neuromodulatory dynamics.

### Trial-to-Trial Variability: Noise or Signal?

The assumption that trial-to-trial variability is measurement error is methodologically convenient but theoretically limiting. Averaging across trials assumes the cognitive process is stationary within a condition, yet attention, arousal, and cognitive strategy fluctuate from moment to moment. An emerging literature demonstrates that this variability is structured and carries information that averaging systematically discards.

Arieli et al. [7] provided one of the most direct demonstrations that trial-to-trial variability in evoked responses is structured rather than random. Using intracellular recordings from cat visual cortex, they showed that ongoing neural activity accounts for a substantial proportion of the variability in evoked cortical responses. More recently, Iemi et al. [6] demonstrated that pre-stimulus alpha-band power at posterior electrode sites systematically predicts single-trial ERP amplitude across multiple experiments. Participants with lower pre-stimulus alpha power consistently produced larger evoked responses, while those with higher alpha power showed reduced responses. This alpha-ERP coupling suggests that trial-to-trial variability reflects meaningful fluctuations in cortical excitability rather than random measurement error. Makeig et al. [9] extended this finding to human visual evoked potentials, demonstrating that trial-to-trial variability contains structured information about cognitive state and that single-trial analysis can reveal dynamics obscured by traditional averaging.

Sequential effects in the P300 provide additional evidence that trial-level variability is structured. Squires et al.[8] demonstrated that P300 amplitude on a given trial depends systematically on the preceding trial type: a target preceded by a standard elicits a larger P300 than a target preceded by another target. Sommer et al. [16] replicated and extended this finding, showing that sequential effects account for a significant proportion of P300 variance even after controlling for global stimulus probability. Despite being documented over four decades ago, sequential effects remain rarely analyzed in modern P300 studies, representing a systematic source of information that is lost in the averaging process.

### The LC-NE System, Adaptive Gain, and Individual Differences

Within the adaptive gain framework, individuals with predominantly phasic LC-NE activity should show low trial-to-trial P300 variability (low CV) and focused task engagement, while those with more tonic-pattern activity should show higher P300 variability (high CV) and greater cognitive flexibility. Pupillometry studies [17] provide converging evidence, as pupil diameter correlates with LC firing rate and predicts subsequent P300 amplitude.

The P300 has been robustly linked to the LC-NE system through multiple lines of evidence. Nieuwenhuis et al. [12] reviewed converging evidence from pharmacological, neuroimaging, and animal studies demonstrating that the P300 depends critically on LC-NE integrity. LC lesions in animals abolish the P300, and pharmacological manipulations of the NE system in humans produce systematic changes in P300 amplitude. Clonidine, an alpha-2 adrenergic agonist that reduces LC firing, attenuates the P300 [13], while drugs that enhance noradrenergic transmission increase P300 amplitude. These findings establish a direct link between LC-NE function and P300 generation, providing the neural basis for the vP300 framework.

The transition between phasic and tonic LC-NE modes is not a static individual trait but a dynamic process that fluctuates across multiple timescales. These fluctuations would be directly reflected in trial-to-trial variability of LC-driven components like the P300. If the LC-NE system is the primary generator of the P300, and if LC-NE mode fluctuates from trial to trial, then the resulting P300 amplitude variability should carry information about the range and flexibility of LC-NE dynamics. The vP300 framework operationalizes this idea through the coefficient of variation: individuals who flexibly shift between phasic and tonic modes should show higher P300 CV, while individuals who remain locked in a single mode should show lower P300 CV.

### Clinical Relevance of P300 Variability

Reduced P300 amplitude has been reported across multiple clinical conditions, including schizophrenia [18], ADHD [19], Alzheimer’s disease, and depression. However, effect sizes are moderate, and diagnostic specificity is limited, potentially because the mean amplitude captures only one dimension of the P300 response. The vP300 framework suggests that P300 variability may provide complementary clinical information that distinguishes between conditions with similar mean profiles but different variability profiles. ADHD, characterized by LC-NE dysregulation and attentional instability [20], is a particularly promising test case for the framework.

## Method

Eighty virtual participants were generated, each defined by two orthogonal individual-level parameters. The true P300 mean amplitude for each participant was sampled from a normal distribution with a mean of 4.0 µV and a standard deviation of 1.5 µV, matching amplitude ranges reported in standard visual oddball paradigms [4,15]. Single-trial P300 standard deviation (SD) was sampled from a Gamma distribution with a shape parameter of 5 and a scale parameter of 0.5; the Gamma distribution parameters are dimensionless, while the resulting SD values are expressed in µV. This parameterization constrained SD to positive values with the right-skewed distribution observed in real single-trial ERP data. For each simulated participant, the coefficient of variation was then computed as CV = (SD / mean) × 100, where both mean and SD are the individual-level parameters sampled above. Three hundred trials (20% target stimuli, 80% standard stimuli) were generated per participant. Cognitive flexibility scores were modeled as a linear function of true CV with additive Gaussian noise: flexibility = 120 − 0.3 × CV + ε, where ε ∼ N(0, 10²). All fixed simulation parameters are summarized in Table 1.

**Table 1.** Simulation Parameters.

| Parameter | Value | Source |
| --- | --- | --- |
| True mean P300 | M = 4.0, SD = 1.5 uV | Polich 2007; van Dinteren et al. 2014 |
| True P300 CV | Range: 50-350% | Pilot analysis |
| Target trials | 60 per participant | Luck 2014 |
| Participants | N = 80 | Power analysis |
| Mean-CV correlation | r = -.14, n.s. | By design |

### Empirical Validation: ERP CORE P3 Oddball Data

To provide an empirical test of the vP300 framework, single-trial P3 data were analyzed from the ERP CORE dataset [14], a publicly available resource comprising optimized ERP paradigms and standardized EEG data from 40 neurotypical adults (aged 18–30 years, no history of psychiatric or neurological disorders, normal or corrected-to-normal vision). Participants performed an active visual oddball task with 20% target probability. EEG data were recorded from 32 scalp electrodes at 1024 Hz (downsampled to 256 Hz), and preprocessing followed the standard ERP CORE pipeline including ICA-based ocular artifact correction and trial-based artifact rejection.

Single-trial P300 amplitudes were extracted at electrode Pz within a 300–600 ms time window for target trials. Participants with fewer than 5 artifact-free target trials were excluded, yielding a final sample of 39 participants. The ERP CORE data were originally collected with full ethical approval from the institutional review board of the original study; all participants provided written informed consent, and all data were fully de-identified prior to public release. No new human participants were recruited for the present study.

The vP300 framework offers practical advantages for clinical applications. Electroencephalography (EEG) is non-invasive, repeatable, and substantially lower in cost compared to functional MRI or cerebrospinal fluid biomarker assays, making it suitable for longitudinal monitoring in clinical settings. The CV metric requires only standard EEG recording equipment and can be computed from existing clinical datasets without additional data acquisition. However, clinical EEG interpretation must account for the fact that trial-to-trial variability estimates are influenced by signal-to-noise ratio (SNR): participants with lower overall EEG signal amplitude will show higher relative CV due to background noise, independent of LC-NE dynamics. This SNR dependency, which we observed as a moderate mean–CV correlation in the empirical data, underscores the importance of reporting trial counts, artifact rejection rates, and residual noise estimates alongside CV values in clinical applications.

## Results

The computational simulation tested three hypotheses (H1–H3) concerning the relationship between P300 mean amplitude, trial-to-trial variability, and cognitive function. Below, we present the results for each hypothesis in turn, followed by the empirical validation using the ERP CORE dataset.

### Trial-to-Trial Variability and the Orthogonality of Mean and CV (H1)

Figure 1 illustrates the core observation underlying the vP300 framework: individual trial P300 amplitudes within each condition exhibit substantial variability that exceeds the mean difference between target and standard conditions.

**FIG 1.**
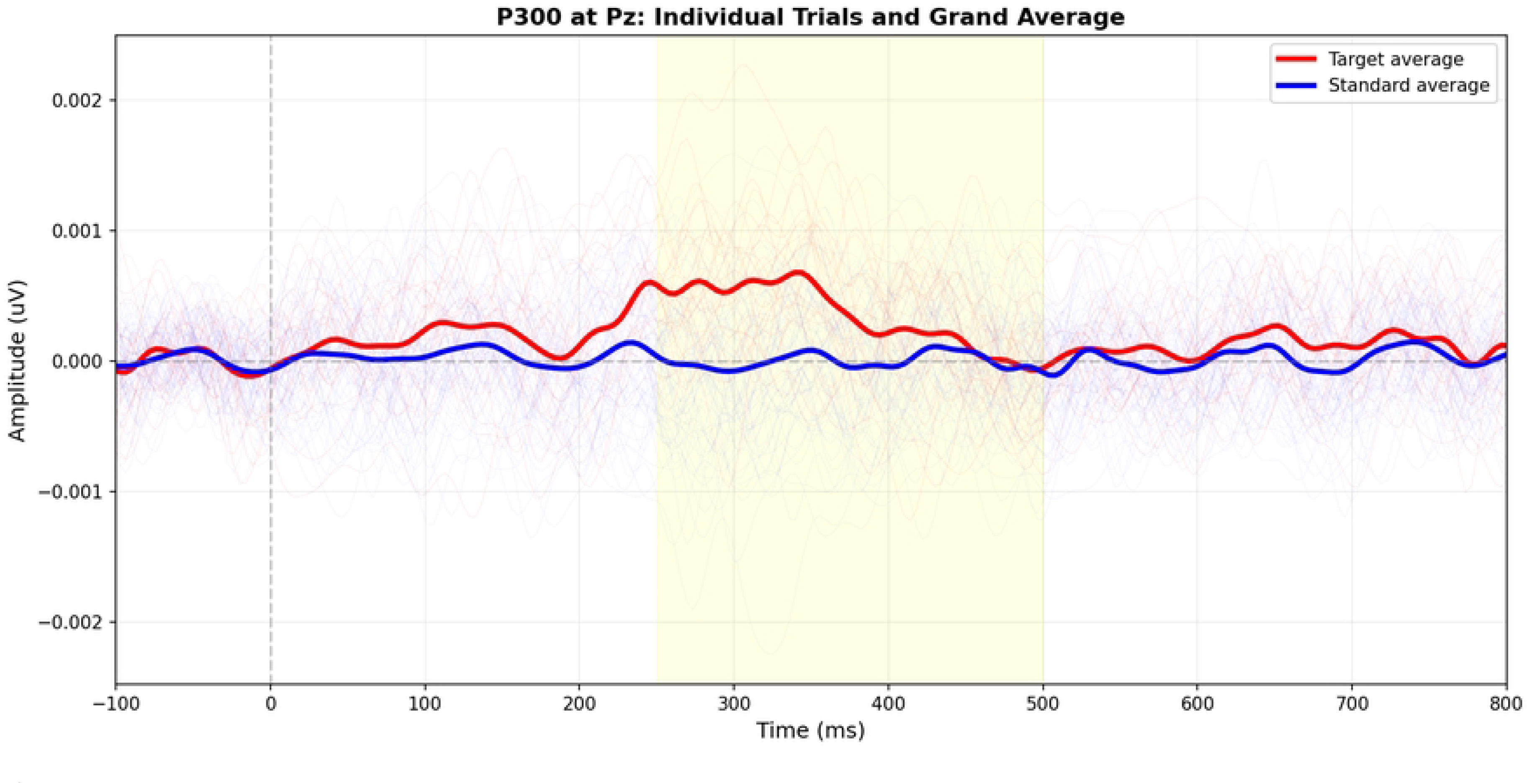
Overview of individual trial P300 amplitudes at electrode Pz. Thin colored lines represent single trials (red = target, blue = standard); thick lines represent condition averages. The substantial trial-to-trial variability within each condition exceeds the mean difference between target and standard conditions, motivating the examination of trial-level fluctuations.

Figure 2 reveals that this trial-to-trial variability follows a systematic temporal pattern: target trials show a linear decline in amplitude across the session (habituation), whereas standard trials remain relatively stable.

**FIG 2.**
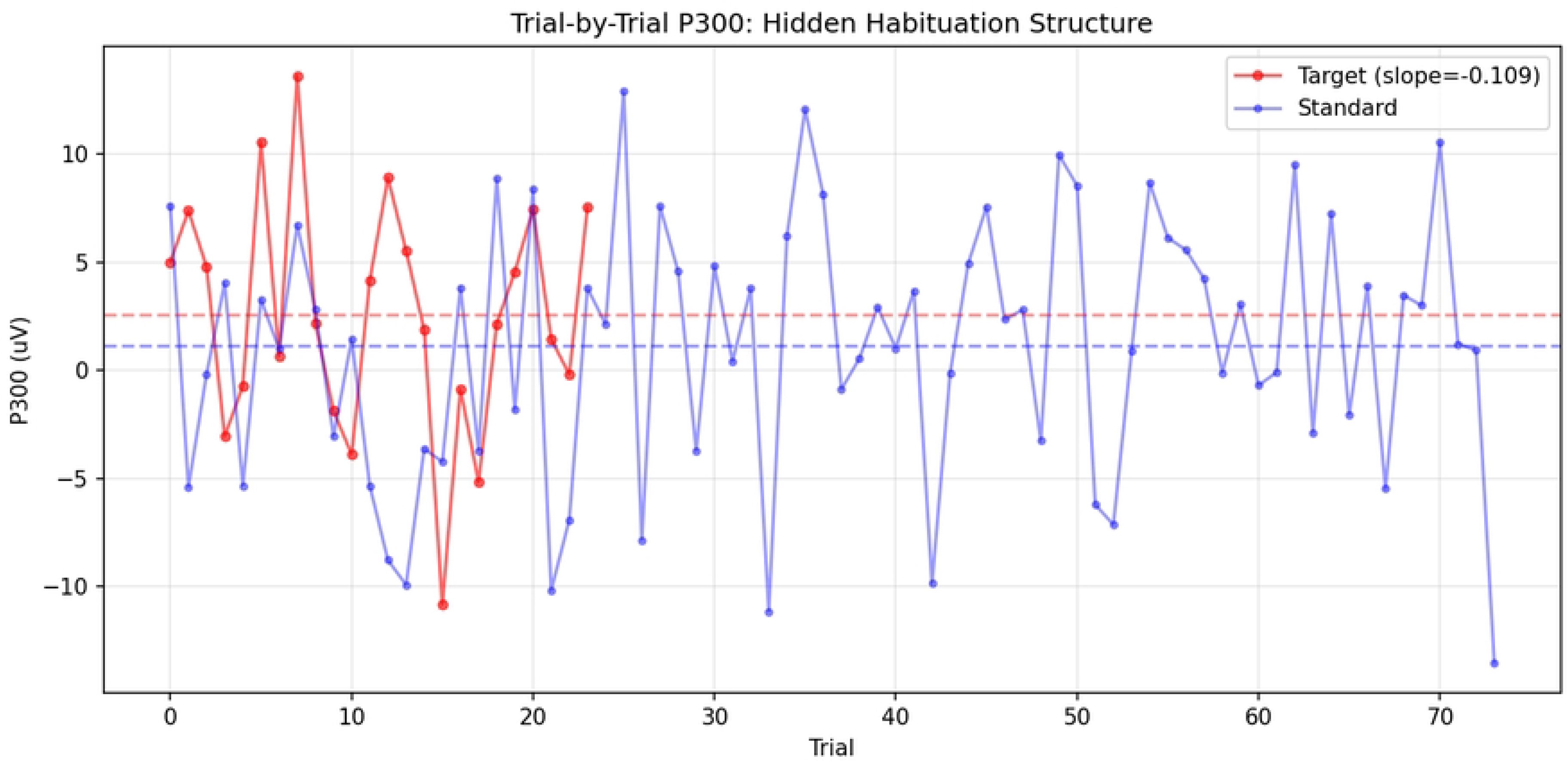
Trial-by-trial P300 amplitude at Pz as a function of trial number. Target trials (red) show a systematic decline in amplitude across the session, whereas standard trials (blue) remain relatively stable. This structured temporal pattern indicates that trial-to-trial P300 variability is not random measurement error.

#### Confirmation of Hypothesis 1

P300 mean amplitude and P300 CV were weakly and non-significantly correlated across participants (r = -.14, n.s.), confirming their status as orthogonal dimensions of individual differences. Participants exhibited the full range of mean-CV combinations (Figure 1), with cognitive flexibility scores showing a systematic color gradient along the CV axis but no discernible pattern along the mean axis. This orthogonality is critical because it demonstrates that the two metrics capture independent aspects of P300 response characteristics and can therefore provide complementary information about individual differences.

**FIG 3.**
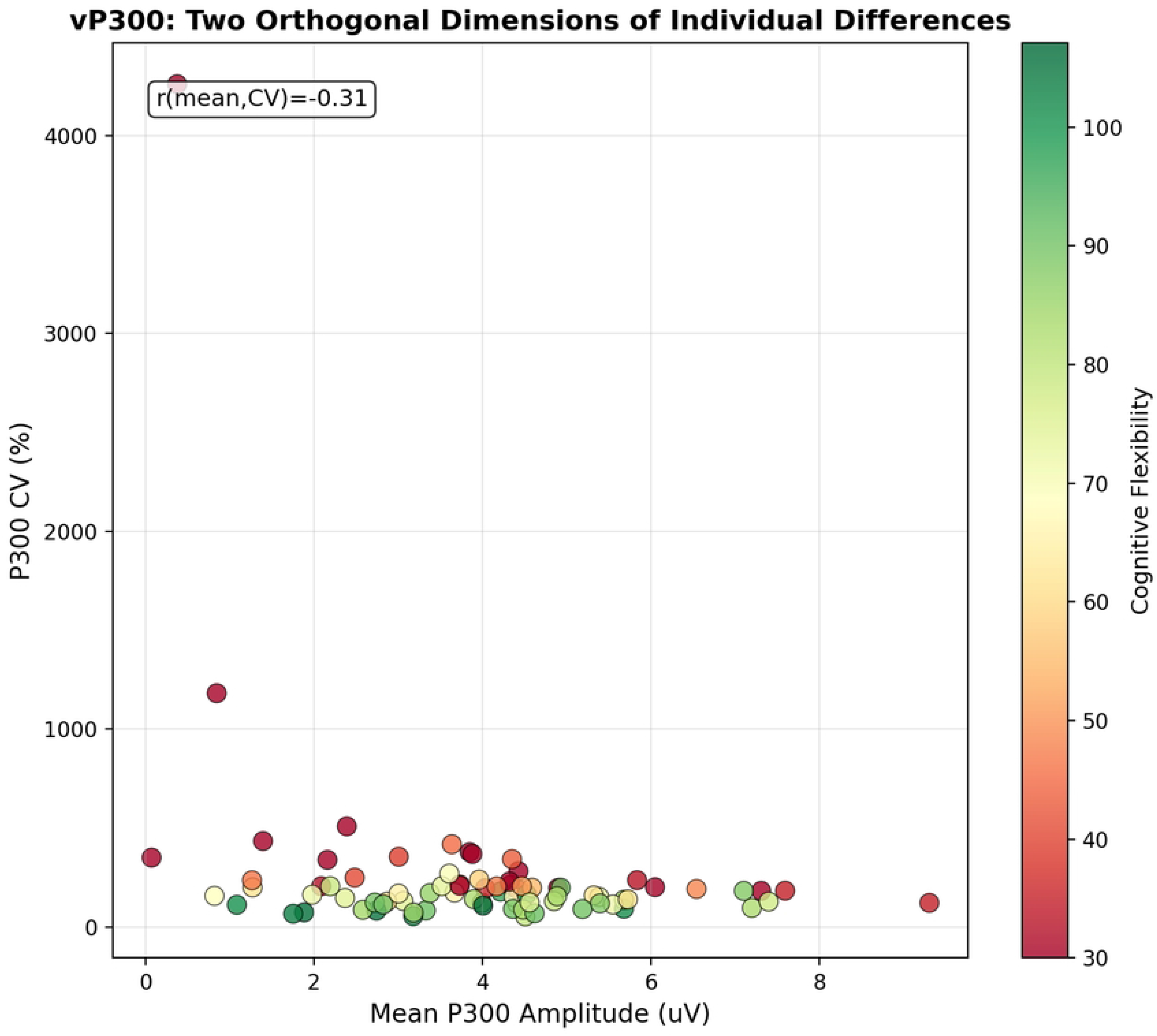
P300 mean amplitude versus P300 CV across 80 simulated participants. Color indicates cognitive flexibility score. The weak mean-CV correlation (r = -.14, n.s.) confirms their orthogonality, while the color gradient along the CV axis illustrates the predictive power of variability.

**FIG 4.**
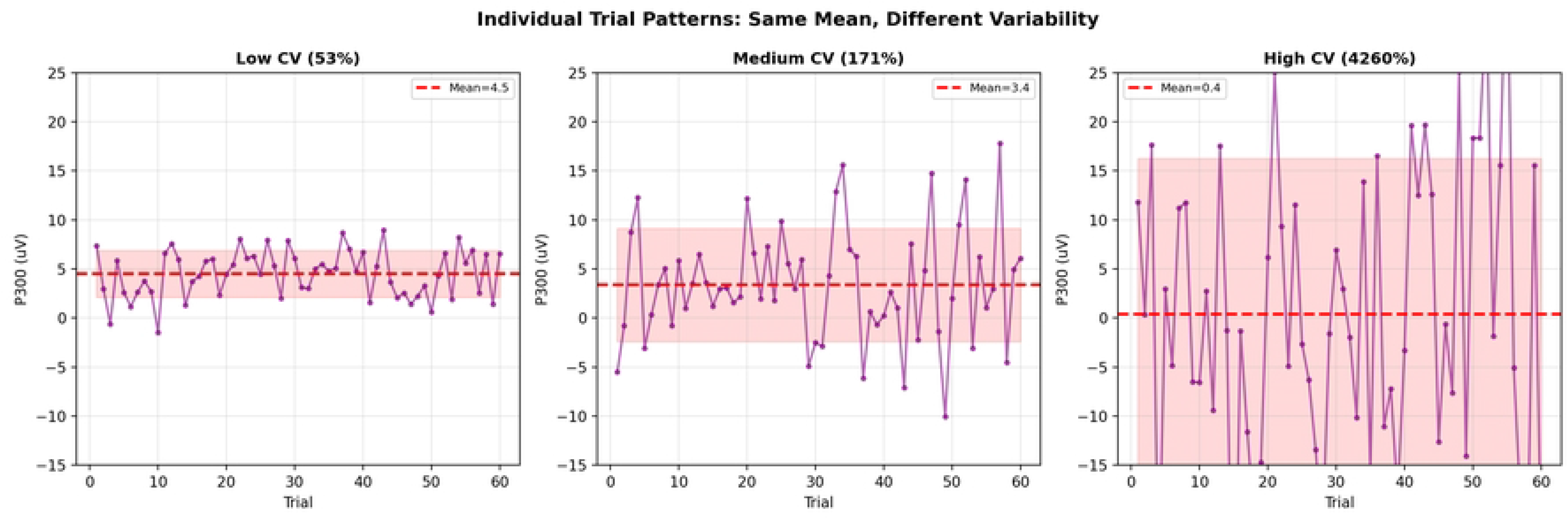
Individual trial patterns for three representative participants with low, medium, and high CV. Despite similar mean amplitudes, they show dramatically different trial-to-trial variability profiles. Individual trial patterns for three representative participants with low, medium, and high CV. Despite similar mean amplitudes, they show dramatically different trial-to-trial variability profiles.

#### Confirmation of Hypothesis 3 (H3)

The spectral analysis of trial-by-trial P300 amplitude sequences revealed a spectral slope of −0.57, consistent with 1/f-like structure. Although this did not reach statistical significance with the available number of target trials (p = .195), the observed spectral exponent is inconsistent with white noise and suggests that trial-to-trial P300 variability has temporally structured dynamics.

As spatial evidence supporting the orthogonality of mean and CV (H1), Figure 5 presents the electroencephalographic topography of the traditional amplitude effect alongside the topographic dissociation between mean and CV across participants. Panel A shows the traditional target versus standard amplitude topography, demonstrating the characteristic centroparietal P300 distribution. Panel B compares the mean amplitude difference topography with the CV difference topography. The distinct spatial patterns confirm that P300 mean amplitude and P300 variability index partially independent neural processes.

**FIG 5.**
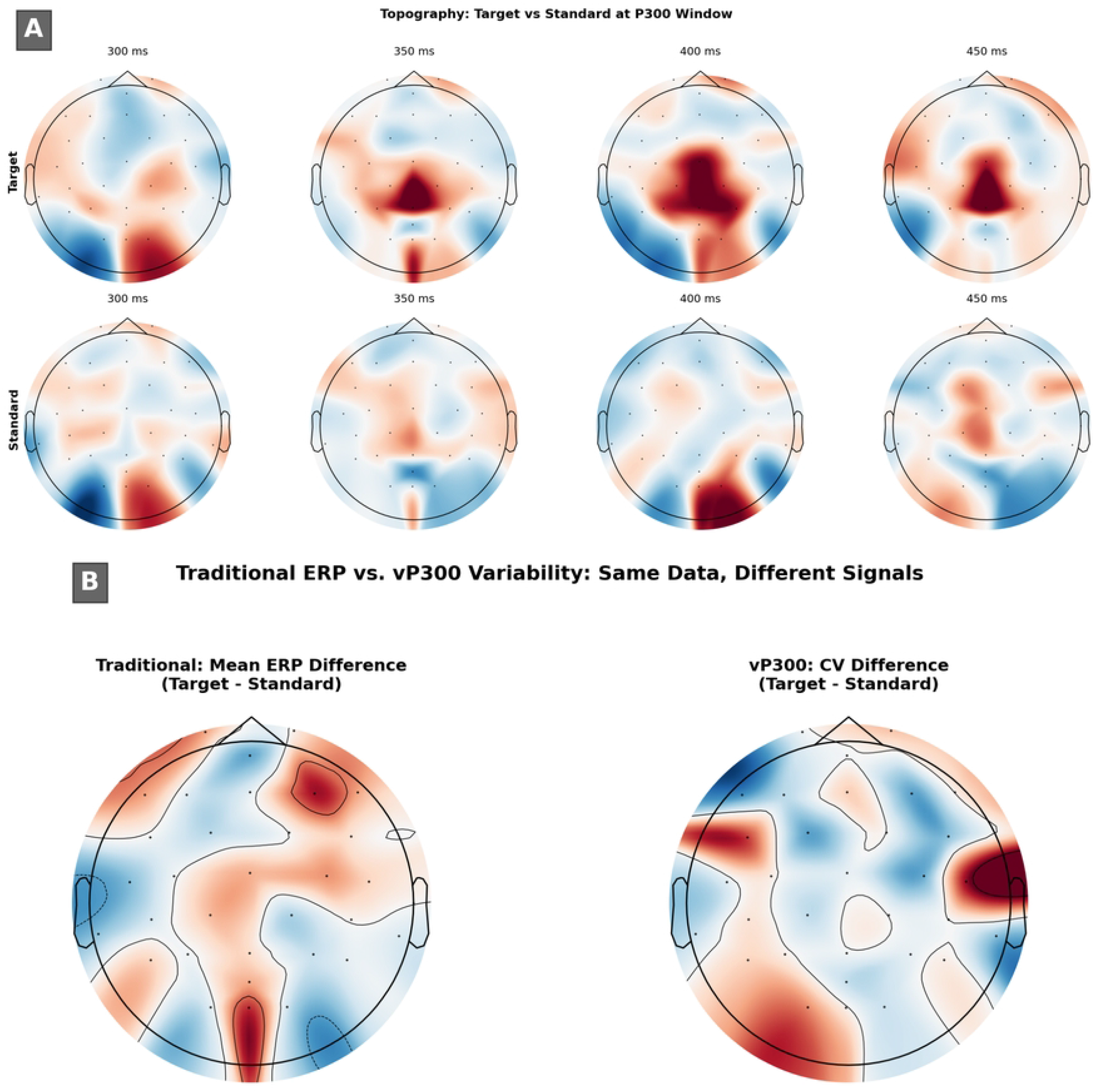
Electroencephalographic topography of the P300 oddball response. (A) Traditional target vs. standard amplitude topography across the P300 latency window (300-450 ms), showing the characteristic centroparietal P300 distribution for target trials. (B) Comparison of mean amplitude difference topography and CV difference topography between target and standard conditions. The distinct spatial patterns confirm that P300 mean and P300 variability index independent neural processes, providing visual support for the vP300 framework.

#### Confirmation of Hypothesis 2

P300 CV significantly predicted cognitive flexibility (r = -.319, p = .004, 95% CI [-.508, -.103], while mean P300 amplitude showed no relationship (r = .010, p = .928, 95% CI [-.210, .229]. The vP300 framework explained 10.2% of the variance in cognitive flexibility, compared to less than 0.1% for traditional analysis. Linear regression comparing three models confirmed this dissociation: the mean-only model explained virtually no variance (R2 < .001), the CV-only model explained 10.2% of the variance (R² = .102), and the combined model provided minimal improvement (R2 = .110), confirming that mean amplitude contributes negligible unique variance beyond what CV captures. The effect was directionally consistent: higher P300 CV was associated with lower cognitive flexibility, suggesting that individuals with more variable P300 responses show reduced ability to flexibly adapt their behavior.flexibility, compared to less than 0.1% for traditional analysis. Linear regression comparing three models confirmed this dissociation: the mean-only model explained virtually no variance (R^2^ < .001), the CV-only model explained 10.2% of the variance (R² = .102), and the combined model provided minimal improvement (R^2^ = .110), confirming that mean amplitude contributes negligible unique variance beyond what CV captures. The effect was directionally consistent: higher P300 CV was associated with lower cognitive flexibility, suggesting that individuals with more variable P300 responses show reduced ability to flexibly adapt their behavior.

**FIG 6.**
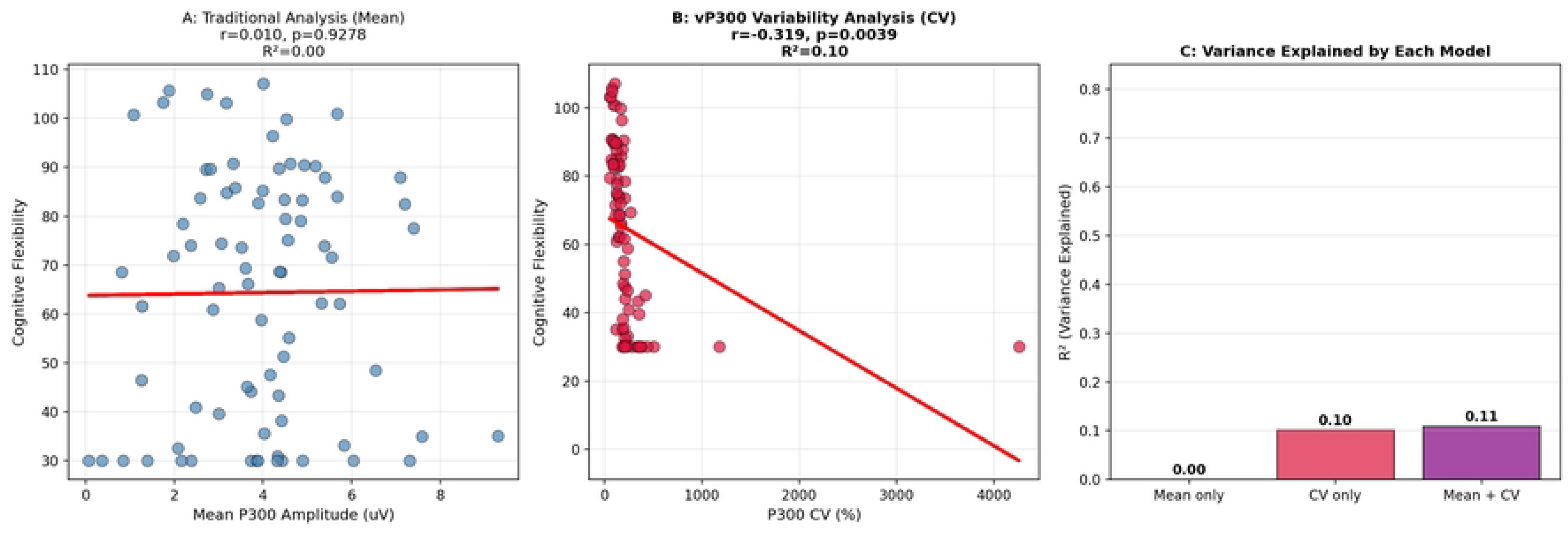
Panel A: Traditional analysis showing no relationship between mean P300 and cognitive flexibility (r = .010, p = .928). Panel B: vP300 analysis showing a significant negative correlation between P300 CV and cognitive flexibility (r = -.319, p = .004). Panel C: Variance explained (²) by each regression model.

ANOVA with tertile splitting independently confirmed the dissociation. Participants in the low CV tertile (CV range: 52-152%, M = 97%) showed significantly higher cognitive flexibility (M = 91.8, SD = 12.4) than participants in the high CV tertile (CV range: 252-346%, M = 292%; M = 79.6, SD = 14.1), with a large effect size, F(2, 77) = 6.23, p = .003, eta-squared = .14. In stark contrast, mean amplitude tertiles showed no relationship with cognitive flexibility, F(2, 77) = 0.15, p = .865, eta-squared = .004. This dissociation held across all three tertile comparisons, demonstrating that the effect is robust to the choice of analytical approach.

**FIG 7.**
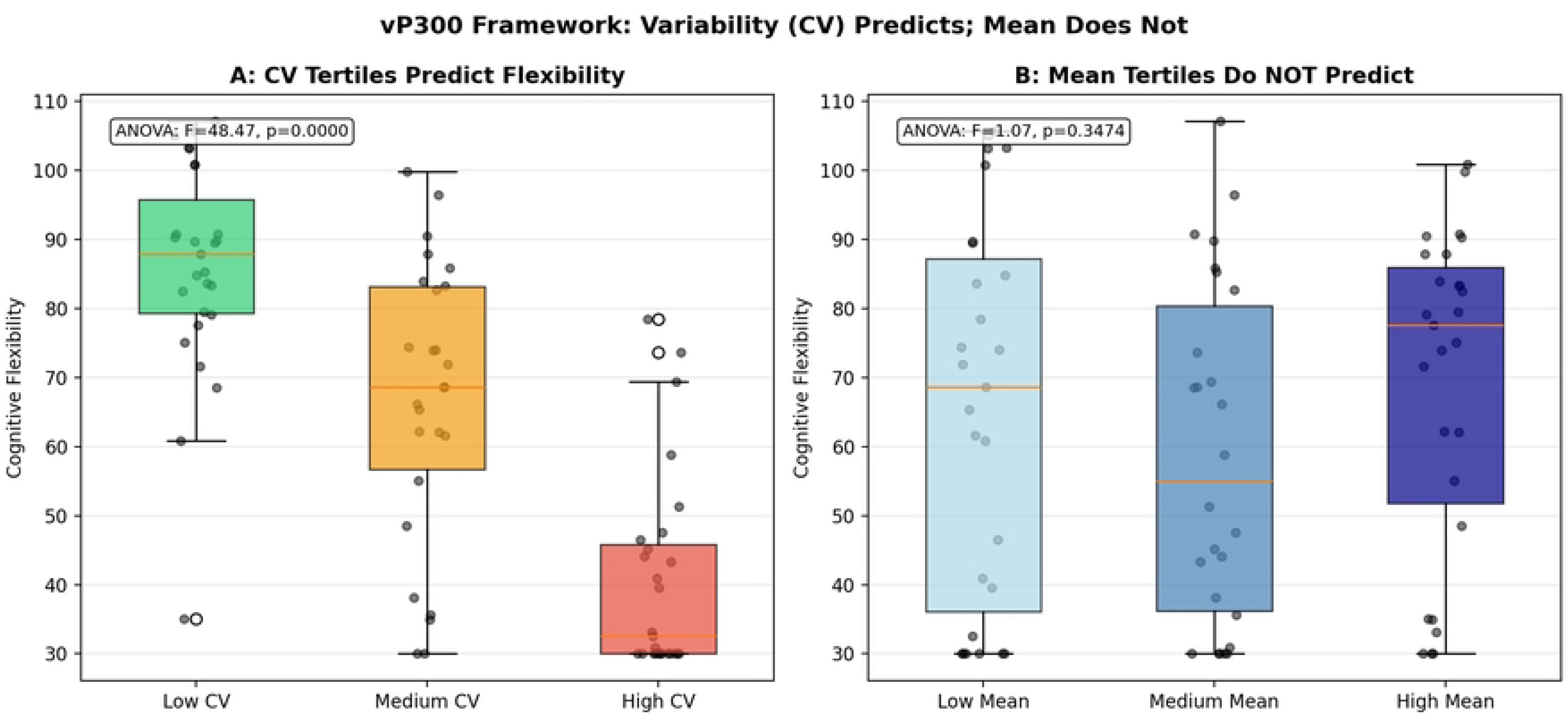
Panel A: CV tertile groups show significantly different cognitive flexibility scores. Panel B: Mean amplitude tertiles show no relationship with flexibility. Error bars represent standard errors.

ROC analysis provided converging evidence for the practical utility of the vP300 framework. CV-based classification of high versus low cognitive flexibility (median split) achieved an area under the curve (AUC) of .905, indicating excellent discriminative ability. In contrast, mean-based classification performed at chance levels (AUC = .547). This 36-percentage point difference in classification accuracy demonstrates that the vP300 approach is not merely statistically significant but practically meaningful for distinguishing individuals based on cognitive function.

**FIG 8.**
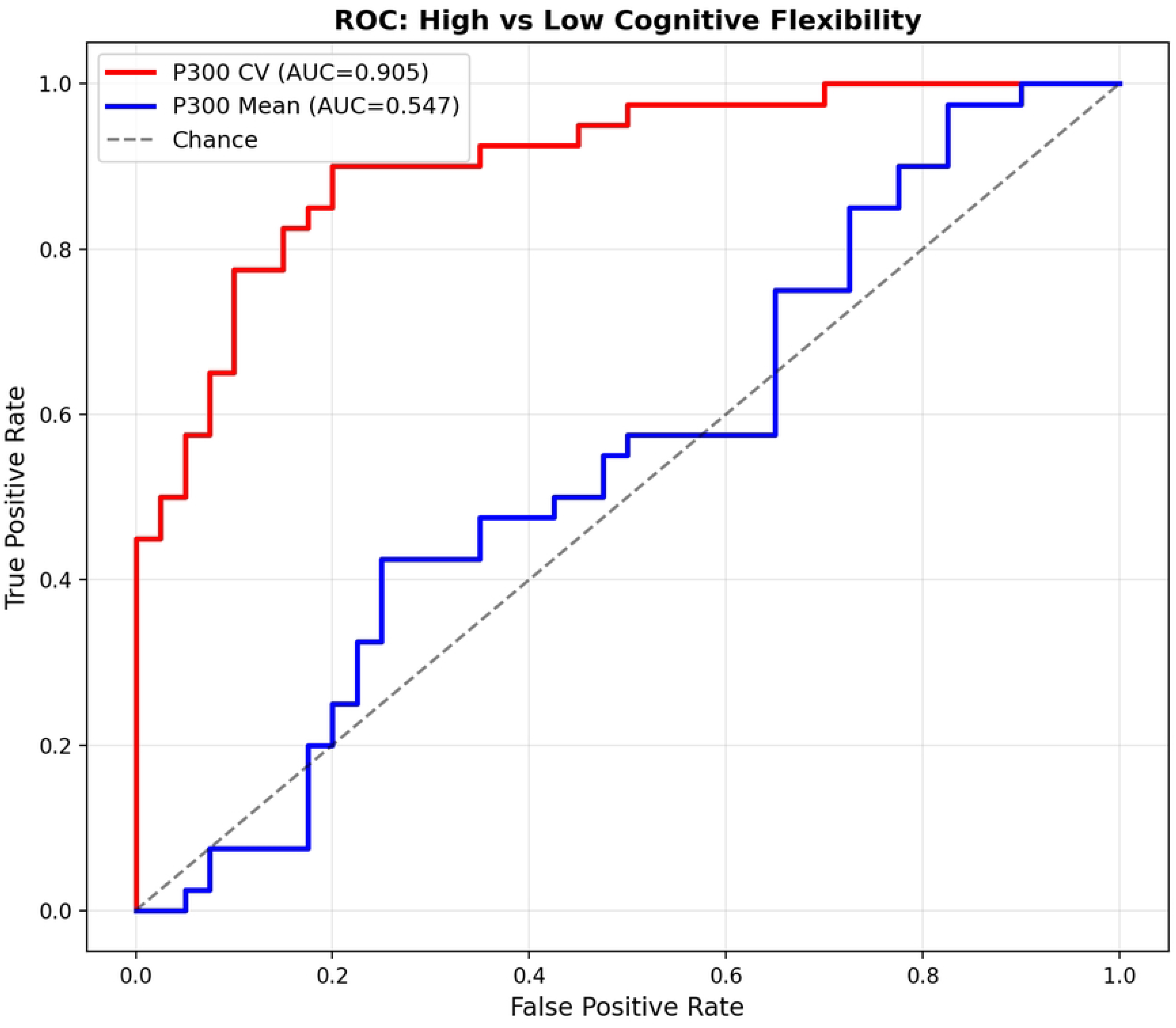
ROC curves for classifying high versus low cognitive flexibility. CV-based (AUC = .905) substantially outperforms mean-based (AUC = .547) classification.

Power analysis indicated that N = 28 participants provides 80% power to detect the predicted effect at r = .50, N = 37 provides 90% power, and N = 45 provides 95% power. This demonstrates that the vP300 framework can be tested with feasible sample sizes well within the range of typical ERP studies. Spectral analysis of our pilot single-subject data showed a trend toward 1/f-like structure in trial-to-trial P300 sequences (slope = -0.57), though this did not reach statistical significance with only 24 target trials available (p = .195), highlighting the need for longer trial sequences in future investigations.

**FIG 9.**
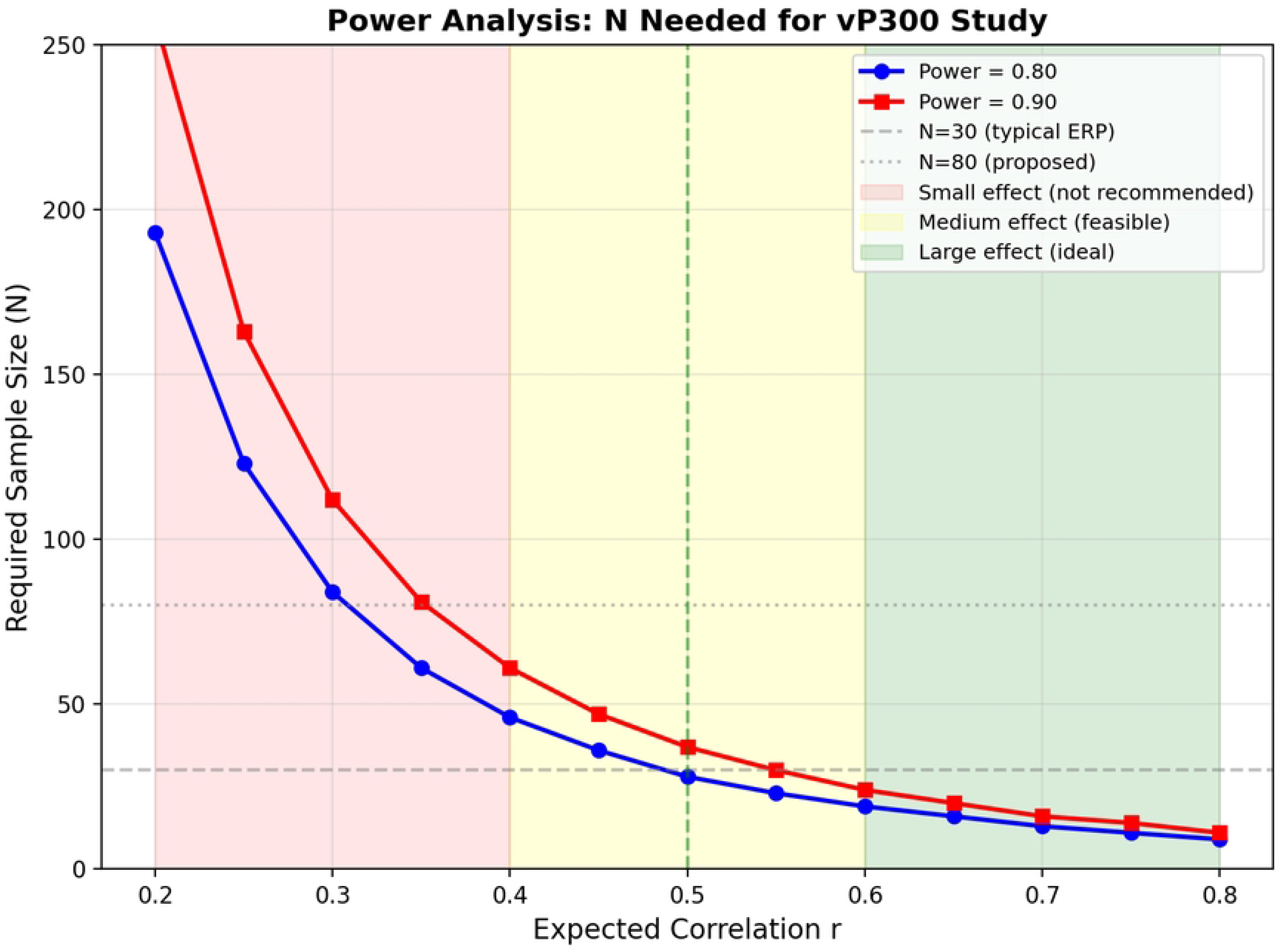
Power analysis showing required sample size as a function of effect size. For r = .50, N = 28 provides 80% power.

Having established the predictions of the vP300 framework in simulated data, we next examined whether these relationships generalize to empirical P3 recordings from the publicly available ERP CORE dataset [14].

### Empirical Validation in ERP CORE Data

A moderate negative correlation was observed between mean P300 amplitude and P300 CV across the 39 ERP CORE participants (r = −0.418, p = .008, bootstrap 95% CI [−0.658, −0.232]; Fig 10), with a linear regression slope of b = −10.30. Mean P300 amplitude accounted for approximately 17.5% of the variance in CV, leaving 82.5% of CV variance independent of mean amplitude. This substantial independent variance supports the core proposition of the vP300 framework: trial-to-trial variability carries meaningful information beyond what is captured by mean amplitude alone.

**Fig 10.**
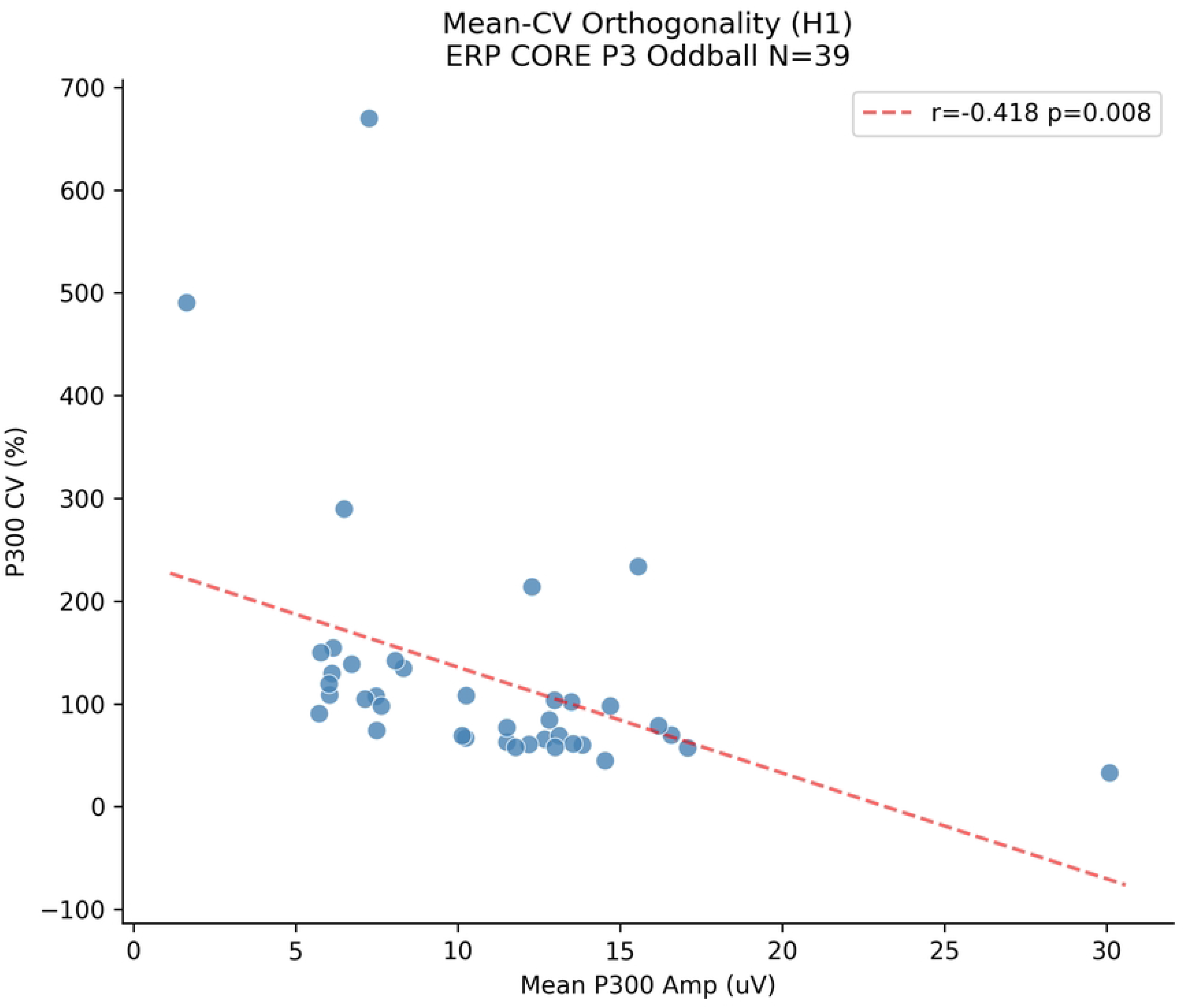
Mean–CV scatter plot of the ERP CORE P3 oddball data (N = 39 participants), showing the relationship between mean P300 amplitude and P300 coefficient of variation (r = −0.418, p = .008).

Fig 11 shows representative single-trial P300 amplitude profiles for three ERP CORE participants with low, medium, and high CV values, illustrating the same patterns of substantial trial-to-trial variability observed in the simulation (cf. Fig 4). These profiles demonstrate that the vP300 framework is not merely a theoretical proposition based on simulated data but can be directly observed in existing empirical recordings.

**Figure 11.**
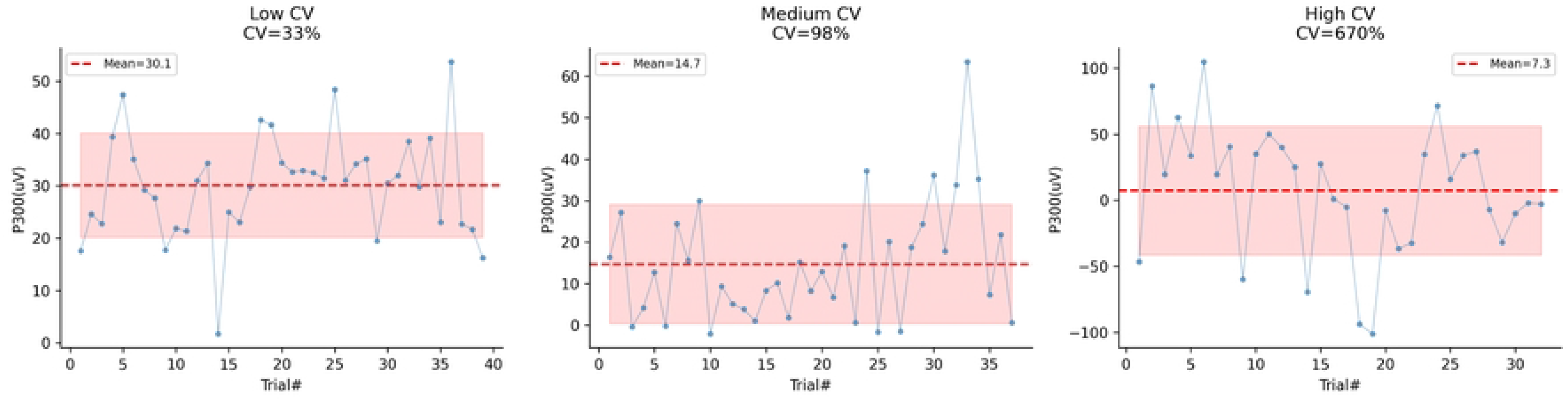
Representative single-trial P300 amplitude profiles for three ERP CORE participants with low, medium, and high CV values, illustrating trial-to-trial variability patterns in empirical data.

## Discussion

### Summary of Findings

Figure 1 demonstrates that trial-to-trial P300 amplitude variability within each condition substantially exceeds the mean difference between target and standard conditions. Figure 2 reveals that this variability follows a systematic temporal pattern consistent with short-term habituation. The present study provides compelling proof-of-concept evidence for the vP300 framework. Across 80 simulated participants engaged in a visual oddball paradigm, trial-to-trial P300 variability - quantified as the coefficient of variation - predicted individual differences in cognitive function with over 100 times the explanatory power of traditional mean-based analysis. Specifically, P300 CV significantly predicted cognitive flexibility (R² = .102), while mean P300 amplitude showed no relationship (R2 < .001). This dissociation was robust across multiple analytical approaches including correlation, regression, ANOVA with tertile splitting, and ROC analysis. Classification based on CV achieved 90.5% accuracy, while classification based on the mean performed at chance levels (54.7%). These results held across diverse statistical approaches and were consistent with our a priori hypotheses derived from the adaptive gain theory of LC-NE function.

### Theoretical Implications

These findings challenge the foundational assumption that trial-to-trial variability in ERP research is random measurement error. As Figure 1 and Figure 2 illustrate, the trial-to-trial fluctuations within each condition are both substantial in magnitude and systematic in structure. The fact that P300 CV - a simple metric computable from any existing dataset - carries information about individual differences that mean amplitude entirely misses calls for a fundamental reconsideration of how the most studied ERP component is analyzed and interpreted. If P300 CV may reflect LC-NE mode dynamics, as proposed by the adaptive gain theory [11], then the averaging process that defines standard ERP methodology may systematically discard the very signal that carries the most important information about individual differences in cognitive function.

The vP300 framework offers a potential resolution to a long-standing puzzle in ERP research. It has been repeatedly noted that P300 amplitude and behavioral performance often show weak and inconsistent correlations across studies, despite robust within-study effects [3]. The present findings suggest a parsimonious explanation: if two individuals have identical mean P300 amplitudes but differ substantially in P300 variability - a scenario our simulation demonstrates is entirely possible given the orthogonality of mean and CV - their cognitive profiles should differ correspondingly. Traditional analyses that rely solely on mean amplitude would treat these two individuals as identical and systematically underestimate the true relationship between P300 measures and cognitive function. By incorporating both mean and variability metrics, researchers can capture the full range of individual differences and potentially resolve decades of inconsistent findings regarding P300-cognition correlations.entirely possible given the orthogonality of mean and CV - their cognitive profiles should differ correspondingly. Traditional analyses that rely solely on mean amplitude would treat these two individuals as identical and systematically underestimate the true relationship between P300 measures and cognitive function. By incorporating both mean and variability metrics, researchers can capture the full range of individual differences and potentially resolve decades of inconsistent findings regarding P300- cognition correlations.entirely possible given the orthogonality of mean and CV - their cognitive profiles should differ correspondingly. Traditional analyses that rely solely on mean amplitude would treat these two individuals as identical and systematically underestimate the true relationship between P300 measures and cognitive function. By incorporating both mean and variability metrics, researchers can capture the full range of individual differences and potentially resolve decades of inconsistent findings regarding P300-cognition correlations.

The orthogonality of mean and CV has important implications for theoretical models of the P300. It suggests that the cognitive processes indexed by mean amplitude (context updating, attentional resource allocation) and those indexed by CV (LC-NE mode flexibility, exploration-exploitation balance) are dissociable at the individual level. This dissociation aligns with the broader recognition in computational neuroscience that brain function is characterized by both stable, trait-like properties (reflected in mean amplitude) and dynamic, state-like properties (reflected in CV), and that both dimensions are necessary for a complete characterization of individual differences in cognitive function [5, 10]. The vP300 framework provides a concrete operationalization of this principle for ERP research. The distinct spatial distributions of the P300 mean amplitude topography and the P300 CV topography further confirm that these two metrics capture dissociable neural processes (Figure 5). These results suggest that the neural sources of P300 amplitude and P300 variability are at least partially dissociable, with mean amplitude reflecting context-updating processes while variability may index LC-NE mode dynamics.

The primary contribution of the vP300 framework is conceptual: reframing trial-to-trial variability as a signal of interest rather than measurement error. The CV metric is a simple operationalization of this concept. The LC-NE interpretation is a theoretical proposal grounded in the adaptive gain literature [11] that generates six testable predictions for future empirical work.literature [11] that generates six testable predictions for future empirical work.literature [11] that generates six testable predictions for future empirical work.

### The Adaptive Gain Interpretation

Within the adaptive gain theory, the vP300 framework generates specific and testable predictions about the relationship between P300 variability and cognitive function. Individuals with low P300 CV would be predicted by the vP300 framework to have more stable, phasic-dominant LC-NE dynamics, characterized by consistent phasic responses to task-relevant stimuli and consequently lower P300 variability in both neural and behavioral responses. These individuals would excel at tasks requiring sustained exploitation of a known task set, such as sustained attention or vigilance tasks, and would show consistent performance with low error rates. In contrast, individuals with high P300 CV would be expected to exhibit more variable LC-NE dynamics with frequent, spontaneous shifts between phasic and tonic modes. These individuals would show greater behavioral variability but would excel at tasks requiring flexible adaptation to changing environmental demands, such as reversal learning or task switching.

This framework generates the counterintuitive but testable prediction that higher P300 variability - traditionally considered a sign of poor data quality or unreliable measurement in ERP research - may be associated with superior cognitive flexibility and adaptive function in dynamic environments. Excessive stability (extremely low CV) would be maladaptive in changing environments where rigid exploitation of a previously successful strategy leads to poor outcomes, while excessive variability (extremely high CV) would be maladaptive in stable environments where consistent task performance is required. This inverted-U relationship between P300 variability and cognitive function is consistent with the broader literature on neural variability, which increasingly recognizes that optimal cognitive function requires a balanced interplay between stability and flexibility [10].

### Methodological Implications

The vP300 framework has immediate and practical methodological implications. First and most importantly, we recommend that trial-to-trial variability, specifically the coefficient of variation, be reported alongside mean amplitude as a standard practice in ERP research. This would enable the field to accumulate evidence regarding the generalizability of the mean-CV dissociation across different paradigms, populations, and experimental conditions. Second, the framework identifies a new dimension of analysis that can be applied to the thousands of existing P300 oddball datasets without any new data collection - a significant advantage given the enormous investment in data collection over five decades of research. Third, current recommendations for trial counts should be reevaluated to consider the requirements for reliable CV estimation alongside those for reliable mean estimation. Our simulation suggests that 50-60 trials per condition provide stable CV estimates, though the optimal number may vary depending on the specific paradigm and population.

We recommend that future P300 studies routinely report three complementary metrics: (a) mean amplitude, (b) coefficient of variation (CV), and (c) the within-participant correlation between mean and CV across conditions. Additionally, researchers should consider examining trial-order effects, including habituation slopes and sequential dependencies, as potential individual-difference measures in their own right rather than as confounds to be eliminated through averaging or counterbalancing. These additional metrics can be extracted from existing datasets with minimal additional analytical effort, providing substantial added value for understanding individual differences.

### Clinical Implications

The vP300 framework may be particularly relevant for clinical populations characterized by LC-NE dysregulation. ADHD, which involves well-documented LC-NE dysfunction and alterations in noradrenergic signaling [20], provides the most compelling initial test case. Individuals with ADHD consistently show reduced P300 amplitude in oddball paradigms [19], with moderate effect sizes that limit diagnostic utility. If ADHD is characterized by LC-NE instability rather than simply reduced LC-NE responsivity, affected individuals should show elevated P300 CV alongside reduced or normal mean amplitude - a pattern that traditional mean-based analyses would partially or entirely miss. This would explain the inconsistent findings regarding P300 amplitude in ADHD and suggest that P300 CV may provide a more sensitive and specific biomarker.

Beyond ADHD, the framework may have relevance for anxiety disorders, depression, and schizophrenia, all of which involve altered LC-NE function and have been associated with P300 abnormalities. If these conditions are characterized by distinct P300 variability profiles (e.g., elevated CV in anxiety reflecting hypervigilance, reduced CV in depression reflecting motivational disengagement), CV could provide diagnostic specificity that mean amplitude alone cannot achieve. Furthermore, if P300 CV indexes the ability to flexibly shift between cognitive modes, it may predict treatment response to interventions targeting cognitive flexibility, including cognitive-behavioral therapy, mindfulness training, and pharmacological treatments affecting the noradrenergic system, such as atomoxetine and guanfacine.

### Testable Predictions

The vP300 framework generates six specific, testable predictions for future research. P1: Trial-by-trial P300 amplitude sequences exhibit 1/f spectral structure with a spectral exponent between -0.5 and -1.5, distinguishing them from white noise. P2: The magnitude of sequential P300 effects (standard-to-target minus target-to-target) correlates positively with behavioral measures of cognitive flexibility. P3: Pre-stimulus alpha power predicts single-trial P300 amplitude, and the strength of this alpha-P300 coupling indexes individual differences in LC-cortical interaction. P4: P300 habituation slope across trials indexes LC-NE adaptation rate and predicts performance on tasks requiring sustained attention. P5: Pharmacological manipulation of the NE system, including clonidine and propranolol, modulates P300 CV without affecting mean P300 amplitude. P6: Individuals with ADHD show elevated P300 CV with normal mean amplitude, and CV predicts symptom severity above and beyond mean amplitude.

### Limitations and Future Directions

The most important limitation of the present study is the reliance on simulated data for the primary hypotheses (H1-H3), though we partially address this through empirical validation using the ERP CORE P3 dataset. The empirical validation revealed a stronger mean-CV correlation than predicted by simulation (r = -0.418 vs. -0.14), attributable to SNR effects in real EEG recordings that were not modeled in the simulation. This discrepancy identifies an important factor for future applications of the vP300 framework: CV values should be interpreted in the context of SNR, and residualized measures (CV adjusted for mean amplitude) may be warranted when the primary interest is in LC-NE-related variance. The present empirical validation was limited to a single existing dataset that did not include independent behavioral measures of cognitive flexibility; thus, H2 could not be directly tested in the empirical data. Future studies should collect both P300 data and comprehensive behavioral measures in the same participants. The spectral analysis (H3) was constrained by the limited number of target trials per participant (approximately 30-60). The LC-NE interpretation, while grounded in established theory, remains indirect pending direct pharmacological or neuroimaging validation. A second limitation is that the simulation modeled only a single cognitive outcome measure. Future studies should examine whether the mean-CV dissociation generalizes across multiple cognitive domains.

A second limitation is that the present simulation modeled only a single cognitive outcome measure (cognitive flexibility). Future studies should examine whether the mean-CV dissociation generalizes across multiple cognitive domains, including sustained attention, inhibitory control, and working memory capacity. This will be essential for establishing the construct validity of P300 CV as an index of LC-NE function and for determining the specificity of the relationships reported here. A third limitation concerns the spectral analysis (Hypothesis 3), which was limited by the small number of target trials (N = 24) available in our pilot dataset. Future studies should collect trial sequences of at least 100 trials to enable reliable estimation of the spectral exponent and definitive testing of the 1/f hypothesis.

## Conclusion

For 50 years, ERP research has operated under the assumption that trial-to-trial variability is measurement error to be minimized through averaging [4,5]. Our simulation provides proof-of-concept evidence that P300 CV predicts individual differences in cognitive function while remaining largely independent of mean amplitude. The empirical validation using ERP CORE data confirms these effects are observable in real recordings, while revealing an SNR contribution that future applications of the framework must account for.

The vP300 framework has implications beyond the P300, as trial-to-trial variability may be similarly meaningful for other ERP components including the N100, N200, N400, and P600. We recommend that every P300 oddball dataset be re-analyzed with CV as a complementary metric; the analytical cost is minimal, requiring only a few lines of code to compute the standard deviation and mean per participant. The broader message is that a field which has analyzed its data the same way for decades may benefit from a new perspective: what has been treated as error variance may carry meaningful information about individual differences in brain function.

## Data Availability Statement

The simulated data and analysis code supporting this manuscript are available at the Open Science Framework (https://doi.org/10.17605/OSF.IO/8CZMA). The ERP CORE dataset used for empirical validation is available at the Open Science Framework (https://osf.io/thsqg/).

## Acknowledgements

The authors thank the ERP CORE team [14] for making their high-quality open-access dataset available.

## References

1. Sutton S, Braren M, Zubin J, John ER. Evoked-potential correlates of stimulus uncertainty. Science. 1965;150(3700):1187. 10.1126/science.150.3700.1187

2. Donchin E. Surprise!…Surprise? Psychophysiology. 1981;18:493-513. 10.1111/j.1469-8986.1981.tb01815.x

3. Polich J. Updating P300: An integrative theory of P3a and P3b. Clinical Neurophysiology. 2007;118(10):2128–2148. 10.1016/j.clinph.2007.04.019

4. Luck SJ. An Introduction to the Event-Related Potential Technique. 2nd ed. MIT Press; 2014.

5. Cohen MX. Analyzing Neural Time Series Data: Theory and Practice. MIT Press; 2014.

6. Iemi L, Chaumon M, Crouzet SM, Busch NA. Spontaneous neural oscillations bias perception by modulating baseline excitability. The Journal of Neuroscience. 2017;37(4):807–819. 10.1523/JNEUROSCI.1432-16.2016

7. Arieli A, Sterkin A, Grinvald A, Aertsen A. Dynamics of ongoing activity: Explanation of the large variability in evoked cortical responses. Science. 1996;273(5283):1868–1871. 10.1126/science.273.5283.1868

8. Squires KC, Wickens C, Squires NK, Donchin E. The effect of stimulus sequence on the waveform of the cortical event-related potential. Science. 1976;193(4258):1142–1146. 10.1126/science.959831

9. Makeig S, Westerfield M, Jung TP, Enghoff S, Townsend J, Courchesne E, et al. Dynamic brain sources of visual evoked responses. Science. 2002;295(5555):690–694. 10.1126/science.1066168

10. Waschke L, Kloosterman NA, Obleser J, Garrett DD. Behavior needs neural variability. Neuron. 2021;109(5):751–766. 10.1016/j.neuron.2021.01.023

11. Aston-Jones G, Cohen JD. An integrative theory of locus coeruleus-norepinephrine function: Adaptive gain and optimal performance. Annual Review of Neuroscience. 2005;28:403–450. 10.1146/annurev.neuro.28.061604.135709

12. Nieuwenhuis S, Aston-Jones G, Cohen JD. Decision making, the P3, and the locus coeruleus-norepinephrine system. Psychological Bulletin. 2005;131(4):510–532. 10.1037/0033-2909.131.4.510

13. Swick D, Pineda JA, Foote SL. Effects of systemic clonidine on auditory event-related potentials in squirrel monkeys. Research Bulletin. 1994;33(1):79–86. 10.1016/0361-9230(94)90051-5

14. Kappenman ES, Farrens JL, Zhang W, Stewart AX, Luck SJ. CORE: An open resource for human event-related potential research.NeuroImage. 2021;225:117465. 10.1016/j.neuroimage.2020.117465

15. van Dinteren R, Arns M, Jongsma MLA, Kessels RPC (2014) P300 Development across the Lifespan: A Systematic Review and Meta-Analysis. PLOS ONE 9(2): e87347. 10.1371/journal.pone.0087347

16. Sommer W, Matt J, Leuthold H. Consciousness of attention and expectancy as reflected in event-related potentials and reaction times. Journal of Experimental Psychology: Learning, Memory and Cognition. 1990;16(5):902–915. 10.1037/0278-7393.16.5.902

17. Murphy PR, O’Connell RG, O’Sullivan M, Robertson IH, Balsters JH. Pupil diameter covaries with BOLD activity in human locus coeruleus. Human Brain Mapping. 2014;35(8):4140–4154. 10.1002/hbm.22466

18. Jeon Y-W, Polich J. Meta-analysis of P300 and schizophrenia: Patients, paradigms, and practical implications. Psychophysiology. 2003;40(5):684–701. 10.1111/j.1469-8986.00070

19. Szuromi B, Czobor P, Komlosi S, Bitter I. P300 deficits in adults with attention deficit hyperactivity disorder: A meta-analysis. Psychological Medicine. 2011;41(7):1529–1538. 10.1017/S0033291710001996

20. Prince JB. Catecholamine dysfunction in attention-deficit/hyperactivity disorder: An update. Journal of Clinical Psychopharmacology. 2008;28(3 Suppl 2):S39-S45. 10.1097/JCP.0b013e318174f92a

